# PIPETS: A statistically informed, gene-annotation agnostic analysis method to study bacterial termination using 3’-end sequencing

**DOI:** 10.1101/2024.03.18.585559

**Authors:** Quinlan Furumo, Michelle Meyer

## Abstract

**Background:** Over the last decade the drop in short-read sequencing costs has allowed experimental techniques utilizing sequencing to address specific biological questions to proliferate, oftentimes outpacing standardized or effiective analysis approaches for the data generated. There are growing amounts of bacterial 3’-end sequencing data, yet there is currently no commonly accepted analysis methodology for this datatype. Most data analysis approaches are somewhat *ad hoc* and, despite the presence of substantial signal within annotated genes, focus on genomic regions outside the annotated genes (e.g. 3’ or 5’ UTRs). Furthermore, the lack of consistent systematic analysis approaches, as well as the absence of genome-wide ground truth data, make it impossible to compare conclusions generated by diffierent labs, using diffierent organisms.

**Results:** We present PIPETS, (**P**oisson **I**dentification of **PE**aks from **T**erm-**S**eq data), an R package available on Bioconductor that provides a novel analysis method for 3’-end sequencing data. PIPETS is a statistically informed, gene-annotation agnostic methodology. Across two diffierent datasets from two diffierent organisms, PIPETS identified significant 3’-end termination signal across a wider range of annotated genomic contexts than existing analysis approaches, suggesting that existing approaches may miss biologically relevant signal. Furthermore, assessment of the previously called 3’-end positions not captured by PIPETS showed that they were uniformly very low coverage.

**Conclusions:** PIPETS provides a broadly applicable platform to explore and analyze 3’-end sequencing data sets from across diffierent organisms. It requires only the 3’-end sequencing data, and is broadly accessible to non-expert users.

## Background

The study of alternative and premature bacterial transcription termination is a rapidly growing field that aims to understand bacterial responses to changes in environmental conditions. The current standard for studying this is 3-prime-end-sequencing (3’-seq)/Term-seq which non- specifically tags the 3’ ends of RNA, allowing for short-read sequencing to map these ends and quantify their relative frequencies. Initial studies using 3’-seq focused on identifying regulatory elements in the 5’UTRs (UnTranslated Regions) of annotated genes by focusing exclusively on signal in the 5’- and 3’-UTRs of these genes [1–3]. This analysis methodology was used or closely mirrored by several studies afterwards, many of which noted an abundance of 3’-seq signal in other genomic regions of the data [4–6]. Other studies created their own custom analysis methods to study 3’-seq data in order to study specific regulatory signals or to add additional sequencing information [7, 8]. Although many different studies have identified that there are biologically relevant 3’-seq signal levels in regions outside of the 3’-UTR of genes, there is not a consensus on an analysis method to best analyze 3’-seq data. As a result, there are no ground truth data sets for the field to measure metrics of success for analysis methods. Furthermore, while analysis scripts are available in many cases, these are typically poorly documented and challenging to implement. Without a standardized 3’-seq analysis method, findings from diffierent studies are impossible to compare short of reanalyzing many large-scale data sets, and it is challenging to contextualize whether findings from diverse organisms are due to diffierences in the biology, the data collection method, or the data analysis performed.

In this work, we introduce **PIPETS** (**P**oisson **I**dentification of **PE**aks from **T**erm-**S**eq data). PIPETS is an R package available on Bioconductor that analyzes mapped data from 3’-end sequencing, which nonspecifically captures transcript termination signal, whether that be from transcript termination, regulatory cleavage, truncation, or other biological processes [9–14]. In contrast to many existing approaches for 3’-seq analysis, PIPETS analyzes all 3’-seq signal, regardless of annotated gene positions, and uses statistically informed methods to identify biologically relevant 3’-end signal. PIPETS provides an analysis platform with many tunable parameters that allows the user to explore their 3’-seq data while also providing statistically grounded results. We demonstrate the use of PIPETS on two existing 3’-seq datasets.

## Implementation

### Overview

The inputs for PIPETS are mapped 3’-seq Bed files that contain at least chromosome, read start, read stop, read quality score, and strand information for each read. Alternatively PIPETS can use Genomic Ranges objects from the <u>GenomicRanges</u> package so long as the input data contains the same information as above [15]. PIPETS begins by filtering out reads that do not have a read quality score equal to or greater than the user defined value (readScoreMinimum parameter). For the data in this study, we noted that using a readScoreMinimum values of 30 resulted in a large reduction in the total reads analyzed (Supplemental Table 1). We chose to use read quality score as an initial thresholding tool to account for the variance in protocols and organisms used in diffierent 3’-seq studies.

This initial trimming step removes reads that do not pass the user defined read quality score threshold, thus improving the overall confidence of the analysis. (Supplemental Table 1). With the quality trimmed reads set by the user inputs, PIPETS calculates the 3’-end read coverage of every genomic position using the terminal position of each read corresponding to the terminal site in the transcript. PIPETS subsequently analyzes this data in three main steps: 1) A sliding window function which uses a Poisson Distribution Test to identify genomic coordinates with significant 3’-end termination read, 2) A condensing step that combines the signal of consecutive significant genomic positions into one termination “peak” and 3) A peak condensing step that combines proximal significant peaks into one signal. These steps ensure that technical or biological error does not conflate noise as significant signal while pruning the total number of individual peaks reported to make analysis manageable (Figure 1).

**Figure 1:**
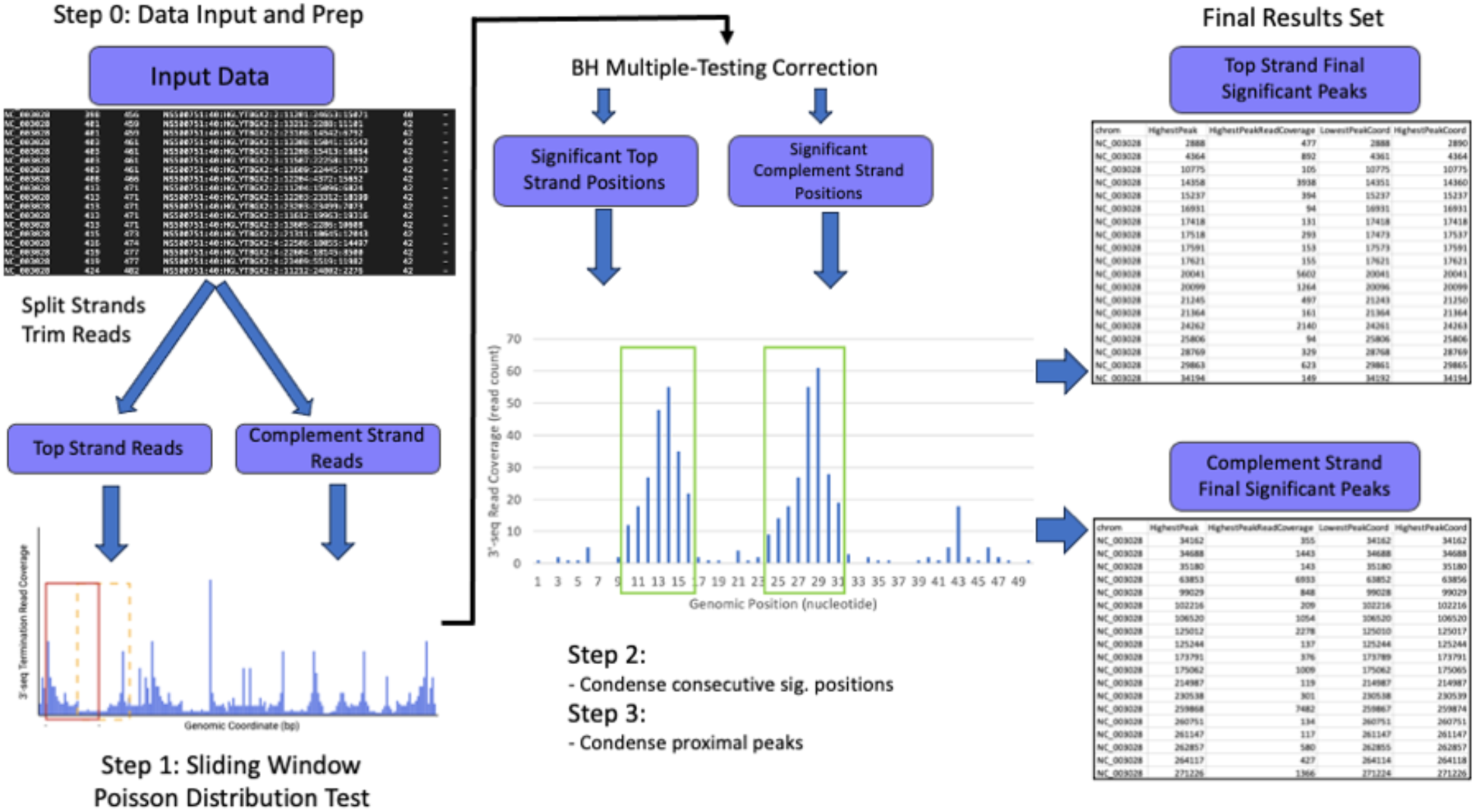
**Workflow of PIPETS**. Before PIPETS performs analysis, it first trims reads based on read quality and separates the passing reads into those that map to the top and complement strands. Then PIPETS analyzes the two strands separately through three steps. First, a sliding window-based Poisson Distribution Test is used to identify genomic coordinates with 3’-seq read coverage that is significantly higher than the surrounding area. The results of this step undergo Benjamini-Hochberg multiple testing correction to control for Type I error. Second, consecutive significant genomic coordinates are combined into individual “peaks”. Third, proximal peaks are combined into individual larger “peaks”.

### Peak identification

One of the inherent challenges associated with analysis of RNA-seq data generally is the large range of read values obtained across different transcripts. Low read coverage does not necessarily indicate a lack of biological significance, as some transcripts have a low basal expression level. However, in the case of 3’-end sequencing, the “noise” created from cleavage of transcripts (either during *in vitro* RNA processing or within the cell) will also be higher for more highly expressed transcripts, potentially much higher than true signal for a lowly expressed transcript. To account for this, PIPETS uses a sliding window approach to ensure that 3’-seq read coverage significance is assigned locally. PIPETS then uses a Poisson Distribution Test to determine if the read coverage of any position within the window is significantly higher than expected from the average read coverage of the window. To minimize the rate of Type I error from the large number of tests performed, PIPETS uses Benjamini-Hochberg multiple testing correction on all significant results. The sliding window approach ensures that genomic positions with read coverage dwarfed by distal highly expressed regions are not overlooked.

Even for input data with millions of reads, a vast majority of genomic coordinates will have very few or no termination reads mapped to them. On average for the Dar data [1], 98.80925% (4163345) of the genomic coordinates had 0 reads with a read quality score of >=30, 99.9499% (4211406) had less than 10 reads, and 99.99305% (4213224) had less than 100 reads. On average for the Warrier data [4], 83.37% (1801494) of the genomic coordinates had 0 reads with a read quality score of >=30, 97.46965% (2106165) had less than 10, and 99.8122% (2156784) had less than 100. (Supplemental Figures 1 & 2). We have higher confidence in genomic coordinates with greater read coverage, however diffierent data sets may have diffiering definitions of “greater read coverage”. To account for such diffierences in the data, we employ a global minimum read coverage cutoff based on the read coverage all mapped reads that pass read quality checks from the data itself rather than an arbitrary predefined minimum read count. Positions with 3’-end read depth below this cutoffi are not tested for significance, speeding analysis and reducing the effect of low-signal noise on analysis. Rather than manually set a specific minimum read-depth prior to analysis, PIPETS determines this global cutoffi using the read coverage of all mapped reads that pass read quality checks in an individual data file.

To determine the global minimum read coverage cutoff, PIPETS first sorts the genomic coordinates from highest to lowest read coverage and identifies the positions that account for a percentage of the total read coverage of the file (defined by threshAdjust, 0.75 by default). The average of these high read coverage values of each file, which are most likely to be derived from biologically significant signal, is the minimum read coverage cutoff for an individual input file.

This results in a global cutoff that is informed by the sequencing depth and signal in the individual experiment rather than an arbitrarily assigned cutoffis. This cutoffi can then be used to filter out noise and reduce run-time by reducing the total number of genomic positions tested.

For the data sets analyzed in this paper, we found that a threshAdjust value of 0.75 provided a cutoffi with values in the hundreds and a runtime that made analysis of many samples quick (Supplemental Table 1). However, for data sets with diffierent read depths or for organisms with potentially diffierent distributions of 3’-seq signal across the genome we anticipate users will modulate threshAdjust to explore their data and best identify peaks at diffiering levels of confidence. A global cutoffi of 5000 3’-seq reads might be an extremely high stringency for some data sets, but could also be required to study 3’-seq data sets with extremely high read depth. To account for possible high read coverage outliers, the parameter highOutlierTrim defines the top percentage (0.01 by default) of these highest read coverage positions that are excluded from the calculation of the global read coverage cutoff. All positions removed this way will still be analyzed by PIPETS, but this helps to ensure that the minimum read coverage cutoffi is not heavily biased by read coverage outliers.

### Peak Condensing Steps

The data structure of a biologically significant bacterial termination site for 3’-seq data resembles a peak, with a position of highest read coverage and flanking positions of lesser read coverage that are still higher than surrounding noise. This signal is the result of several factors, namely imperfect bacterial termination [16, 17], RNA degradation [18], and technical noise/bias from the library preparation and sequencing itself [19]. Each genomic position from the significant results of The Poisson Distribution test in Step One of PIPETS is unlikely to be an independent termination signal; many of these positions are adjacent to others and together they are likely to represent one favored site of termination (Supplemental Figure 3). To better reflect the biological context of these data, the significant genomic positions identified in Step One undergo two condensing steps: 1) Combine consecutive significant positions into single termination peaks, and 2) Combine proximal termination peaks to abbreviate total results volume (Figure 1). Both condensing steps preserve the total range of data combined, so that the information about the breadth of peaks is not lost. However, by reducing the total results pool, these steps help to make the results of PIPETS easier to digest and study.

Step Two of PIPETS identifies significant positions that are within 2 nucleotides of each other. These significant positions are then combined into a single termination “peak” which contains the full range of consecutive positions and uses the highest read coverage position within for downstream analysis. Step Three of PIPETS combines proximal termination peaks within 20 nucleotides of each other as termination signals so close to each other are likely not independent sites of transcript termination as both Rho-dependent and independent termination require RNAP pausing (reviewed in [17, 20]), which typically occurs on strong hairpin structures of at least 7-8 base pairs in length yielding a minimum distance of 15-20 nucleotides between termination sites [20–22].

### PIPETS Parameters

PIPETS relies on a number of parameters, the values of which initially were generated in an *ad hoc* manner. These include the size of the sliding window (slidingWindowSize), the distance the sliding window moves with each step (slidingWindowMovementDistance), the p-value used for the Poisson distribution test and Benjamini-Hochberg multiple testing correction (user_pValue), the proportion of data used to determine the global minimum read cutoffi (threshAdjust), the proportion of data considered to be high outliers in the calculation of the global minimum read cutoffi (highOutlierTrim), the distance between adjacent positions considered to be a single peak during the first peak condensing step (adjacentPeakDistance), and the maximum distance between adjacent peaks condensed in the second peak condensing step (peakCondensingDistance).

We established the default values of these parameters using *S. pneumoniae* 3’-seq data generated in our lab [4] with the intention of identifying as many results as possible that we were confident had the potential to be biologically significant (which may be distinct from statistically significantly greater than background). To systematically do so, we assessed the impact that changes to values of each parameter had on results sets. We first tested the sliding window portion of the analysis. By default, the sliding window spans 51 genomic positions, moving 25 nucleotides with each step. Alternative sizes and movement distances were tested, but did not confer any notable change to analysis. We selected this size because it provides a small enough snapshot of genomic regional context for analysis while also containing enough total genomic coordinates to ensure that the Poisson test conducted within the range is sufficiently populated.

For the peak condensing steps, we found that, outside of extremely high or low values, these parameters had little impact on the total number of results and did not heavily impact the read depth cutoffis or the total count of significant results (Supplemental Figures 4,5). However, changes to the global minimum read coverage cutoffi substantially impact the number of peaks returned by PIPETS. This value is determined from the input data by two PIPETS parameters: threshAdjust and highOutlierTrim, and testing revealed that minor adjustments to these values can greatly affiect the distribution of read coverage values that are assessed. Altering the threshAdjust and highOutlierTrim values changes the strictness of PIPETS. In datasets with very high read depths or datasets from very long genomes, we recommend increasing the strictness of PIPETS’ noise filtering by lowering the value of threshAdjust (from 75% to values from 50% - 60%). This will reduce the effect of global increases to basal noise levels and ensure that PIPETS does not consistently misidentify lower read coverage events as significant 3’-seq signal.

Alternatively, increasing the value of threshAdjust (from 75% to values from 80% - 95%) will make PIPETS less strict and is useful when studying data sets with low read coverage or data from smaller genomes. In the event where a small portion of the genomic positions in the data account for a disproportionately large percentage of the total read coverage for a data set (for example in samples with poor rRNA depletion), increasing the value of highOutlierTrim (from 0.01 to values from 0.025 – 0.05) can reduce the bias imparted by these outliers. highOutlierTrim is the more sensitive parameter and its values may be more variable from dataset to dataset. Although threshAdjust and highOutlierTrim had the largest impact on analysis for the data sets used in this study, we invite users to change the other parameters for their data to better explore the transcription termination profile of diffierent data sets from diffierent organisms.

### Comparison Methodology

In this work we compared the results of PIPETS with the results of the original 3’-seq analysis method that was used for data sets from two organisms. Our goals were to ensure that PIPETS was identifying significant 3’-seq termination signal across all regions of the input data and that PIPETS maintained the ability to detect the high read coverage 3’-seq termination signal found by the previous analysis methods. First, we examined *S. pneumoniae* 3’-seq data from two conditions (no-drug-control time 0 and no-drug-control time 30) that were generated in our lab (available at NCBI SRA: SRP136114) [4]. We ran this data at default parameters (threshAdjust =0.75 and highOutlierTrim = 0.01) before categorizing the results as 3’-UTR (less than or equal to 150 bp downstream of the STOP codon with respect to strand), 5’-UTR (less than or equal to 450 bp upstream of the START codon with respect to strand), coding region (inside of the coding region of a gene), 3’-UTR & Coding region (less than or equal to 150 bp downstream of the stop codon of one gene with respect to strand and inside the coding region of a different gene), and intergenic (none of the other categories). To stay consistent with the previous analysis method [1], we chose for 5’-UTR signal to override 3’-UTR in our reported metrics. In Figure 2A we represent the originally reported peaks all as 3’ ends as originally reported. We used the TIGR4 genome annotation (refseq: NC_003028.3) to assign the above categories. When comparing the results of the original Warrier method and PIPETS, we classified overlapping results when two peaks reported from the different analyses were within 5bp of each other (Figure 2A, Figure 2B).

**Figure 2:**
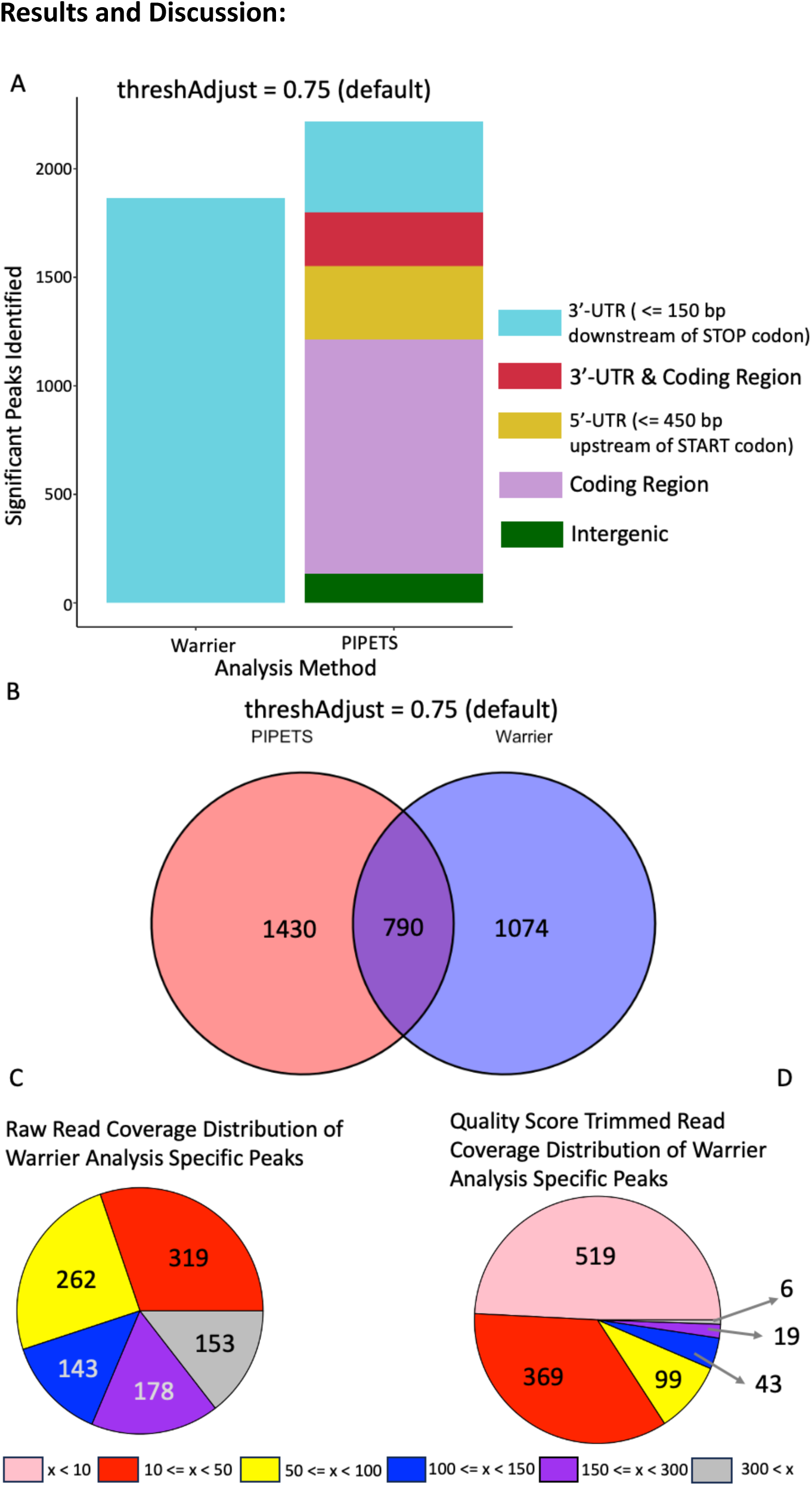
PIPETS analysis of Warrier 3’-seq data. We analyzed the Warrier 3’-seq data on PIPETS using default parameters (threshAdjust = 0.75, highOutlierTrim = 0.01). A) Compared to the 1864 significant sites reported in the 3’-UTR of annotated genes from the original results [4], PIPETS identified 2220 significant peaks. 419 of these peaks were in of the 3’-UTR (within 150 bp of the stop codon) of annotated genes, while the other 1801 were found in other genomic contexts that are unique to PIPETS. B) When comparing the two results sets, PIPETS and the Warrier analysis identified 790 shared peaks. C) When using the raw 3’-seq read coverage of the remaining 1074 results that were unique to the Warrier analysis, only 153 had 3’-seq read coverage greater than 300. D) When using only reads with a read quality score of 30 or higher, only 6 of the Warrier analysis specific results had read coverage greater than 300.

When studying the Warrier results specific peaks, we calculated the read quality trimmed read coverage values based on the combined 3’-seq read coverage from the ND0 and ND30 samples combined (Figure 2D).

Second, we assessed *B. subtilis* data from the first paper to present 3’-seq as a method (Bioproject PRJEB12568)[1]. This study had 9 experimental conditions with three replicates for all conditions but one, resulting in 26 total files. We analyzed all of these files individually and followed the Dar protocol of sub setting the results to only those that were present in all three replicates of a given condition. We then performed the same categorization and comparative results overlapping as above. We used the NCBI <u>AL009126</u> genome annotation for *B. subtilis* in our results categorization. For this part of the work, we ran PIPETS twice, once with default parameters (threshAdjust = 0.75 and highOutlierTrim = 0.01) and once with only one changed parameter (threshAdjust = 0.85).

## Results and Discussion

We first tested PIPETS on *Streptococcus pneumoniae* data generated by our own lab [4]. Previous analysis of this data was performed using an analysis method that mirrored that of the first 3’-seq paper [1]. We developed PIPETS to analyze this data to better capture 3’-seq signal outside of just the 3’-UTR of annotated genes and thus we wanted to compare PIPETS’ results with the original method. The original method identified 1864 termination sites across three conditions, and as a result of the analysis parameters many of these identified sites had very low read coverages. PIPETS, when run with default parameters, identified 2220 significant peaks between just two of these conditions (no-drug-control samples at 0 and 30 minutes). Of these 2220 peaks, 136 were in intergenic regions, 340 were in the 5’-UTR of a gene, 1077 were inside coding regions, 419 were in the 3’-UTR of a gene, and 248 that were present in the 3’-UTR of one gene and inside the coding region of a different adjacent gene (Figure 1A, Supplementary Data). PIPETS identified 1801 significant peaks outside of 3’-UTR regions, all of which would not have been considered by the original analysis method.

Although PIPETS was successful in identifying 3’-seq signal outside of 3’-UTR regions, we still wanted to identify how successful PIPETS was at identifying the results from the original analysis. Of the 1864 sites from the original analysis, PIPETS identified 790 (Figure 2B). It was expected that not all of the sites identified by the original analysis would be identified as significant by PIPETS due to the substantially more lax method of peak identification in the original method [4]. Of the 1055 unique remaining sites of the original analysis, 902 had 3’-seq read coverage values less than 300 (Figure 2C). We wanted to identify the cause of PIPETS “missing” 153 of the results from the original method that had high read coverage values. An important distinction in analysis between PIPETS and the original method is the read quality trimming that PIPETS performed. We compared the raw read coverage and quality score trimmed read coverage at the locations identified by the original analysis and saw that only 6 of the results from the original analysis had more than 300 reads that passed read quality cutoffis (Figure 2D). We further investigated the specific peak identified by the original analysis with the highest 3’-seq read coverage not identified by PIPETS. This position was identified as significant by PIPETS, however, it was placed into a larger 3’-seq peak with a diffierent highest read coverage coordinate that may have been outside the distance parameters in the original analysis.

The largest disparity between the PIPETS results and the Warrier results was the dramatic increase in significant 3’-seq peaks identified outside of 3’-UTR’s of genes for the PIPETS results. Among these results, there are clear instances of 3’-seq signal inside of coding regions of genes, and there are several instances of 3’-seq peaks that could potentially function as premature terminators in genes. Observing the read pile up data for at these sites in IGV, it is clear that there are 3’-seq read buildup events that are of 50% or greater magnitude in relation to the read buildup at the expected termination site at the end of the gene. These premature sites identified by PIPETS strongly indicate that the *ptsP* and SP_RS10700 (Figure 3A and 3B respectively) genes have biologically relevant premature termination sites. These events are 172 and 207 bp past the start codon of their respective genes, and would not be captured by existing 3’-seq analysis methods. Additionally, PIPETS identified a novel un-identified site in the data between the *rpsG* and *fusA* genes (Figure 3C). PIPETS identified significant 3’-seq peaks at the annotated termination sites of both genes (indicated with black arrows) but it also identified significant amounts of 3’-seq signal in a region between both genes. IGV read buildups and PIPETS read coverage outputs suggest that there is biologically significant event causing substantial 3’-end coverage in this area. This signal too, would have been entirely missed by the original analysis method since the signal does not fall within the close constraints of being within 150 bp of the stop codon of a gene.

**Figure 3:**
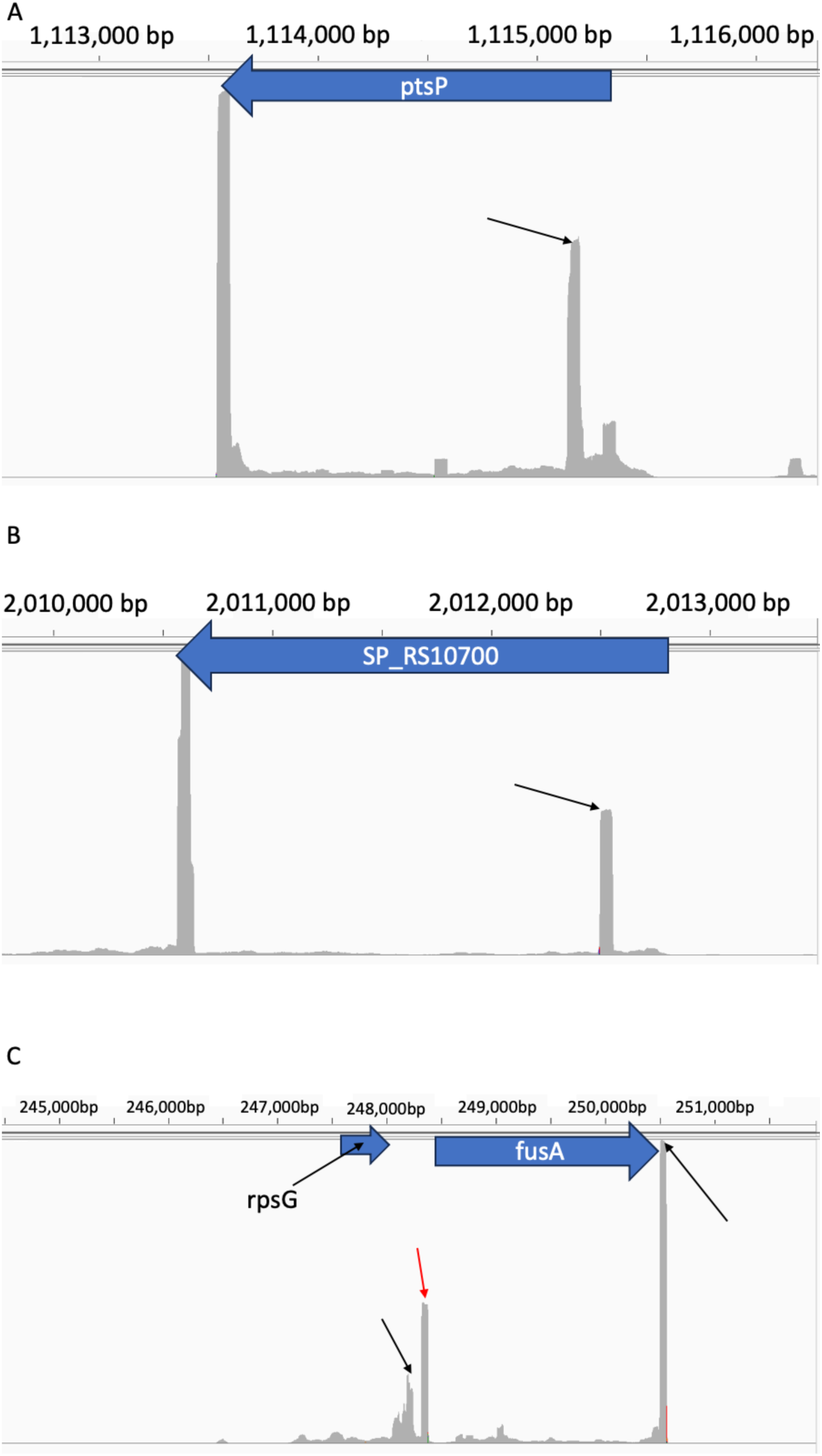
Representative selection of PIPETS specific results from Warrier data. A) PIPETS uniquely identified a significant 3’-seq peak inside of the *ptsP* gene 172 bp after the start codon. IGV display of read pileup data shows that this peak has read coverage within the same range of the likely true terminator signal in the 3’- UTR. B) PIPETS uniquely identified a 3’-seq peak inside of the SP_RS10700 gene. This peak has roughly half of the read pileup of the likely terminator signal, but its read pileup in comparison to the surrounding area indicates likely biological relevance. C) PIPETS identified significant 3’-seq signal in the expected 3’-UTRs of *rpsG* and *fusA* genes (black arrows), but also identified a significant peak in the intergenic region between these two genes (red arrow). This PIPETS specific peak has read pileup values higher than the *rpsG* terminator signal and half as much as the potential *fusA* terminator suggesting biological significance.

We next tested PIPETS on *Bacillus subtilis* data from the first paper to implement 3’-end sequencing as a method of studying bacterial termination [1]. The sequencing depth of the original data was low relative to modern sequencing standards, with no replicate sequencing file with more than 3 million aligned reads. Their analysis methodology focused only on the position of highest termination signal in the 5’-UTR and 3’-UTR of annotated genes, did not consider reads strand specifically, and excluded all other data from analysis. We were curious whether a more systematic analysis method would alter conclusions of the original work. From their combined data, Dar et al. identified 1443 total termination positions that passed their minimum read thresholds and were in the 5’-UTR or 3’-UTR of genes of interest.

We first ran PIPETS on the default parameters and compared that results set to the results of the Dar method. With “threshAdjust” set to 0.75, PIPETS identified 362 significant peaks: 5 were in intergenic regions, 154 were in the 5’-UTR of genes, 14 were inside coding regions, 170 were in the 3’-UTR of a gene, and 19 were present in the 3’-UTR of one gene and inside the coding region of a diffierent adjacent gene (Figure 4A). PIPETS identified 240 of the 1443 sites identified by the original method (Figure 4B), and while more than half of these remaining sites had 3’-seq read coverage values less than 50 (Figure 4C), we still wanted to ensure that PIPETS was properly able to study this data even if most of the identified results had low 3’-seq read coverages.

**Figure 4:**
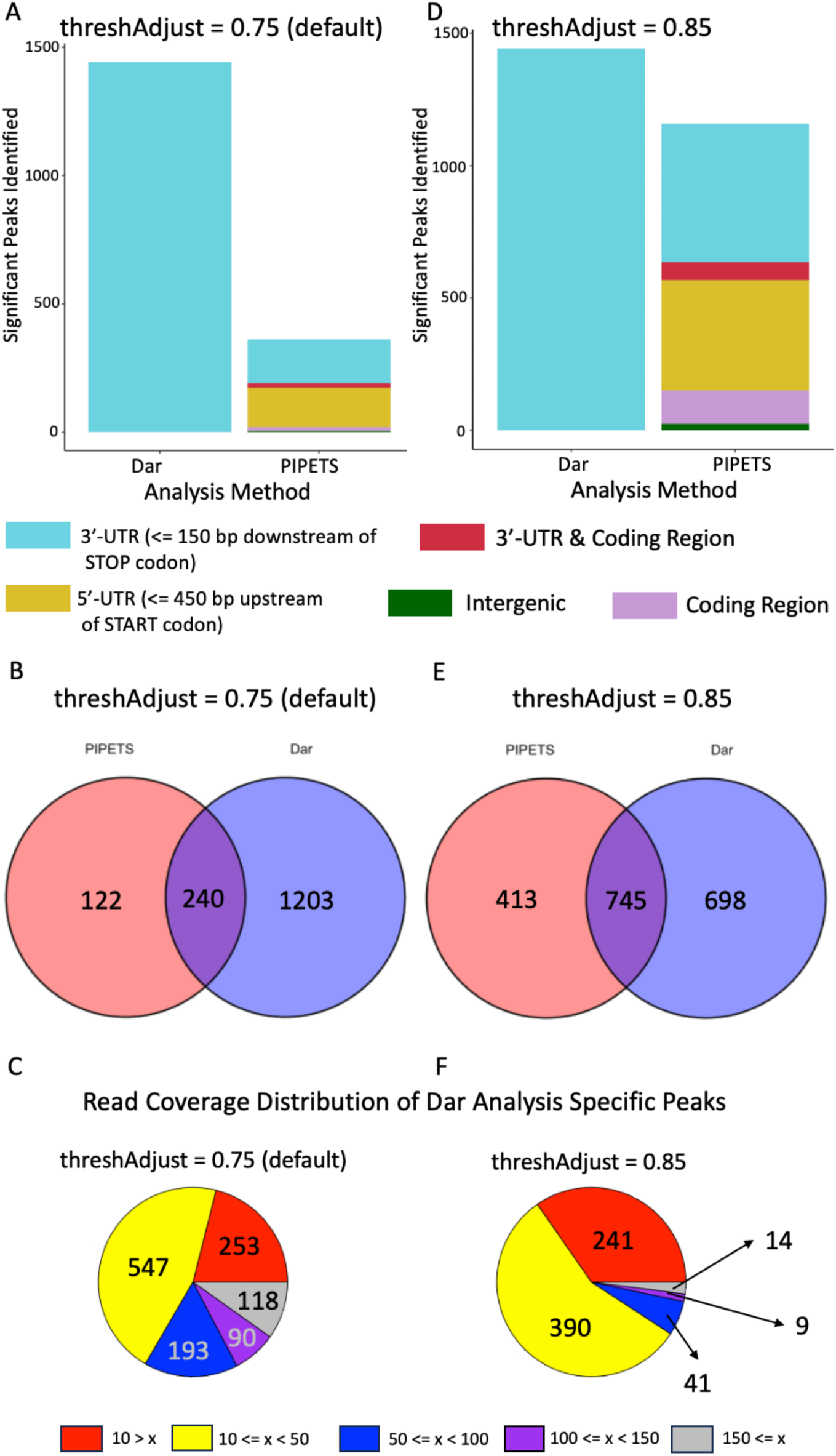
PIPETS analysis of Dar data. A) Using default parameters on the Dar data, PIPETS identified 362 significant peaks, primarily in 3’-UTR and 5’-UTR regions. 3’-UTR peaks for Dar data are as originally reported [1] B) PIPETS identified 240 of the results from the original Dar data and had 122 unique results. C) Of the remaining 1203 Dar analysis specific results, 1085 had 3’-seq read coverage values less than 150 and 800 had read coverage less than 50, which can be too low to distinguish from noise. D) To increase the number of peaks that PIPETS identified, we reduced the strictness of PIPETS by increasing threshAdjust to 0.85. This resulted in PIPETS identifying 1158 significant peaks across a broader range of genomic regions. E) PIPETS identified 745 of the peaks found by the original analysis with these parameter changes, and identified 413 unique 3’-seq peaks. F) Of the 698 Dar analysis specific peaks, only 14 had read coverage values greater than 150. The parameter change of the PIPETS analysis increased its sensitivity to identifying lower read coverage 3’-seq signal.

In order to improve PIPETS’ performance for this data, we changed the “threshAdjust” parameter from 0.75 to 0.85 to make PIPETS more sensitive to lower 3’-seq read coverages. With this change, PIPETS identified 1158 significant peaks, 25 were in intergenic regions, 417 were in the 5’-UTR of a gene, 126 were inside coding regions, 522 were in the 3’-UTR of a gene, and 68 that were present in the 3’-UTR of one gene and inside the coding region of a diffierent adjacent gene (Figure 4D). Notably, this singular change to PIPETS increased the number of significant peaks from 362 to 1158, and also helped to greatly increase the number of significant peaks identified inside of coding regions as well as peaks present in the 3’-UTR of one gene and inside the coding region of a different adjacent gene. When compared to the original results set, PIPETS was now able to identify 745 of the original method’s peaks (Figure 4E). With only 64 of the remaining unidentified peaks having 3’-seq read coverage greater than 50 (Figure 4F), we were confident that PIPETS was appropriately analyzing this data set and identifying most 3’-seq signal that originated from biologically significant events, yet not imposing arbitrary constraints on which data are included in the analysis.

We were curious if the dramatic increase in significant peaks identified by PIPETS upon parameter changes had any recognizable pattern in the data. We tested if the change in number of peaks found in coding regions had any preferential changes. A large increase in the number of genic peaks identified within a small number of highly expressed genes would potentially indicate an increase in calling noisy 3’-end coverage within high coverage areas as legitimate peaks. The results run with “threshAdjust” = 0.75 had very few genes with any significant peak identified inside, and there were no genes that had more than one peak inside for the top strand and no genes with more than 2 peaks for the complement strand (Figure 5). When PIPETS was run with “threshAdjust” = 0.85, while there was a large increase in the total number of genes with significant peaks inside, there was not a proportional increase in the number of genes with more than one significant peak inside (Figure 5). While there is no expected value of potential alternative termination sites inside of any given coding region, we would not expect there to be many genes with several termination sites (greater than 5) due to potential negative overcomplexity. The increased sensitivity PIPETS (“threshAdjust” = 0.85) rarely identified more than 3 peaks within a single gene, suggesting that the parameter change increases PIPETS sensitivity to identifying signal, without mistakenly over-identifying noise within gene coding regions as significant results.

Among the significant peaks identified by PIPETS that were not identified in the Dar method, there are many instances of clear 3’-ends occurring within genes. We identified two examples of PIPETS specific significant peaks that corresponded with clearly biologically relevant sites for genes in the *B. subtilis* genome. When visualized with IGV, read buildups at the annotated ends of the *rlmKB* and *recQ* genes (Figure 6A and 6B respectively) drops offi dramatically after the position identified by PIPETS as a significant 3’-seq read coverage peak. This indicates a standard termination signal where the large buildup of 3’-seq reads at the site identified by PIPETS provides insight to a termination event that was undetected by the Dar analysis. We also noted that the Dar method largely ignored signal associated with various non-coding sequence annotations, including tRNAs. While tRNA’s are not often considered in analyses of this nature, they still make up a significant portion of the total 3’-seq read depth for these files. We highlight here that PIPETS identified significant 3’-seq peaks for a set of 14 tRNA’s in a ∼2000 bp region that directly correspond with IGV signal in the same positions (Figure 6C). While such sites are accessible via reanalysis of the raw data, this omission during initial analysis points toward the challenges of comparing diffierent datasets, generated by diffierent labs, using diffierent organisms, analyzed with *ad hoc* tools, in order to gain broader insight into how 3’-end generation diffiers across bacterial species.

**Figure 5:**
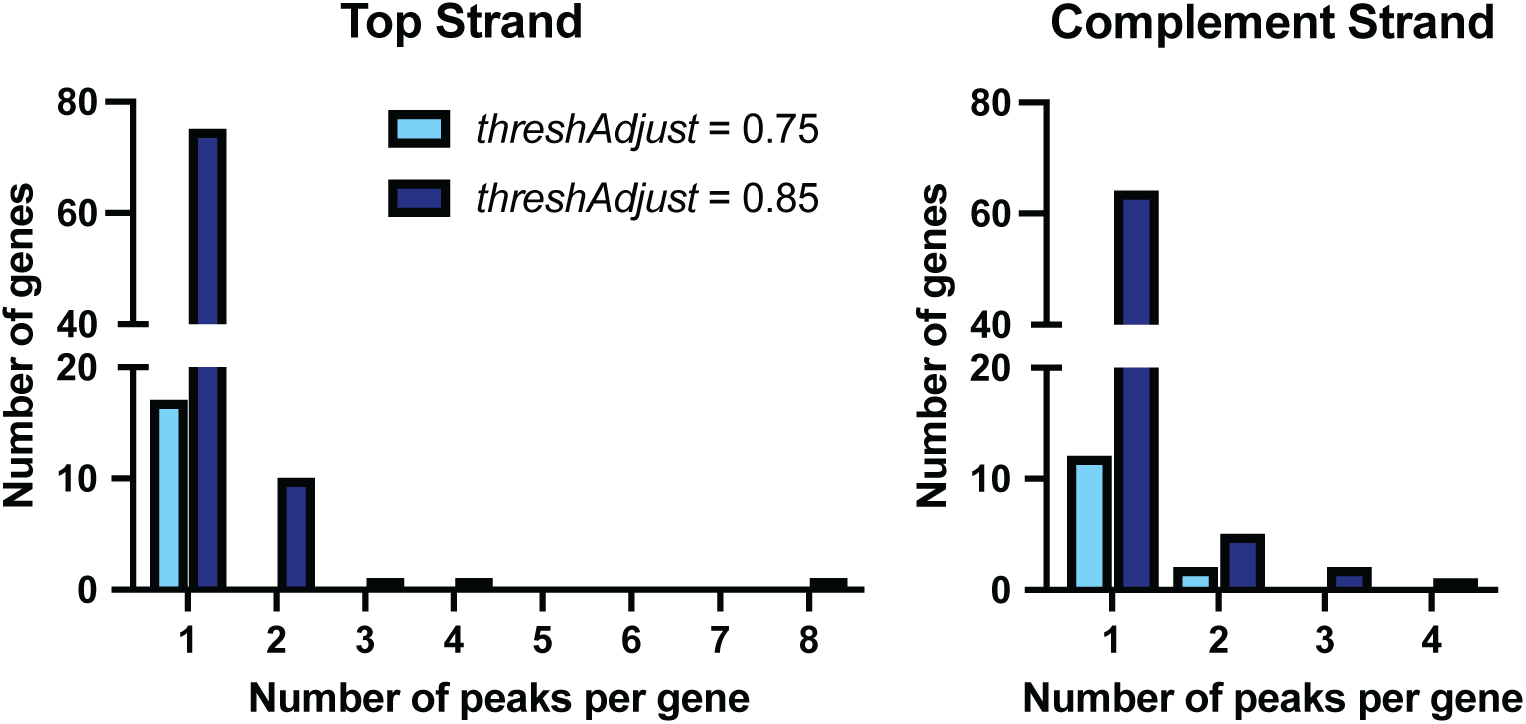
Peaks per gene changes based on threshAdjust for Dar Data. While the number of total significant peaks identified by PIPETS increases with change to threshAdjust, the number of genes with multiple peaks per gene does not disproportionately increase. This means that PIPETS is increasingly sensitive to 3’-seq signal inside of more total genes without increasing the rate of misidentifying excess peaks from noise inside of genes.

**Figure 6:**
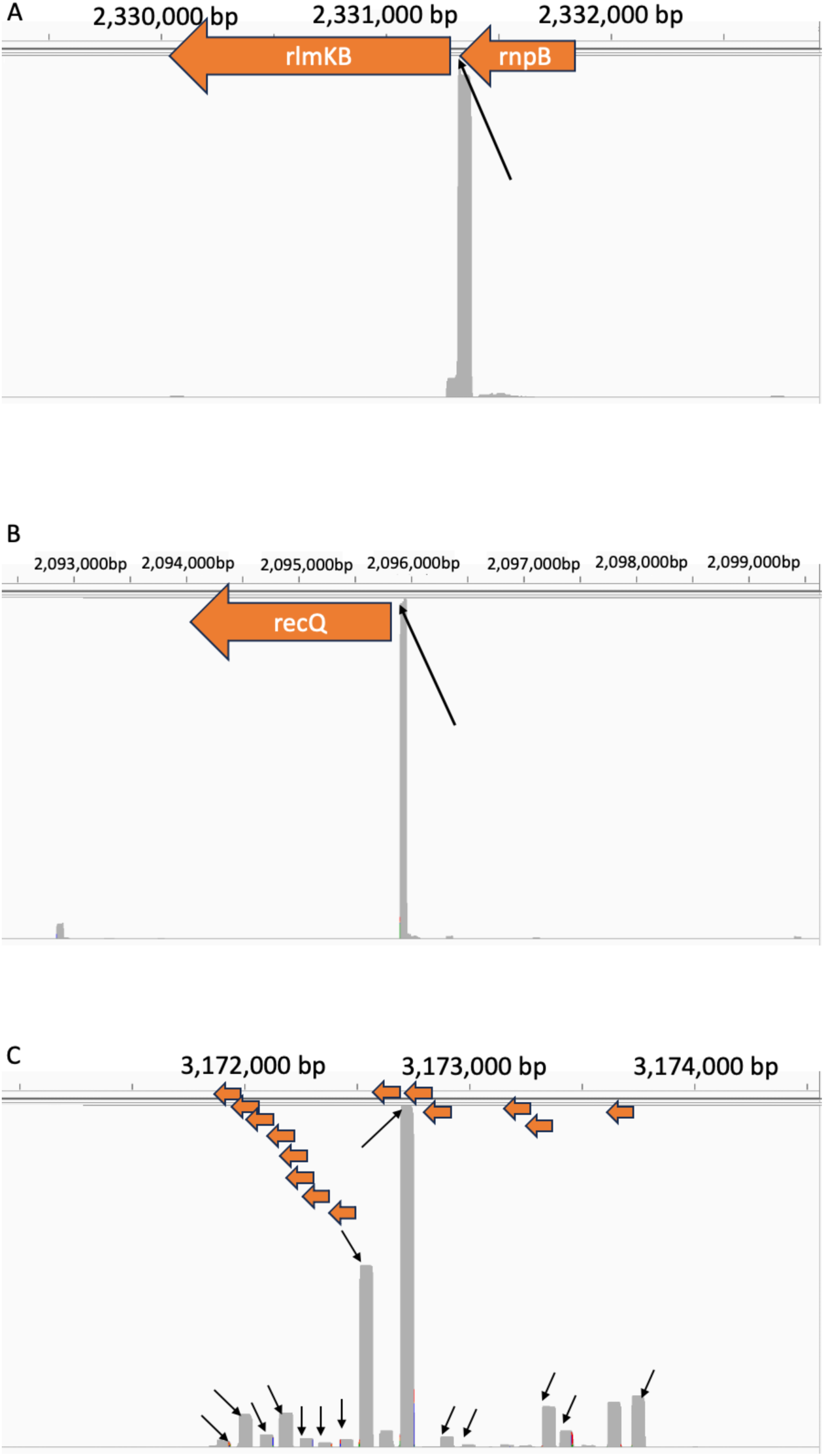
Representative selection of PIPETS specific results from Dar data. A) PIPETS uniquely identified a 3’- seq peak that is inside of the *rnpB* gene and in the 5’-UTR of the *rlmKB* gene. The positioning of this peak could allow it to function as a regulating signal for both genes. B) PIPETS uniquely identified a significant peak in the 5’-UTR of the *recQ* gene. The IGV displayed read buildup indicates that this peak has dramatically higher read coverage than the surrounding region, suggesting biological relevance. C) PIPETS uniquely identified 14 significant 3’-seq peaks from 14 tRNA’s in a ∼2000 bp region. Each of the peaks indicated by arrows were not identified by the Dar analysis, and while tRNA’s are often not the focus 3’-seq studies, these peaks account for all of the 3’-seq signal in these areas and the magnitude of these peaks suggests biological relevance.

## Conclusions

The number of studies utilizing 3’-seq is rapidly increasing across diverse sets of organisms, yet there is not an easy to download, standardized analysis method currently available. Here we present PIPETS, a new 3’-seq analysis method that identifies statistically significant 3’-seq signal in a gene annotation agnostic fashion. When compared to existing 3’-seq analysis methodologies, PIPETS identified many significant 3’-seq signals outside of the 3’-UTR of genes, without sacrificing the ability to identify most of the highly expressed 3’-seq sites in the 3’-UTRs. The tunable parameters of PIPETS allow for the adjustment of the sensitivity of the analysis, which allows PIPETS to analyze data of varying read depths as well as account for biological diffierences between diffierent species.

When studying the data presented here, we noted a large proportion of the significant identified peaks were present outside the 3’-UTR of annotated genes. While altering the *ad hoc* definitions of 5’- and 3’-UTR to be more inclusive of coding regions may capture more of these peaks, such definitions may vary across diverse organisms and ultimately remove biologically relevant sites from analysis based on arbitrary constraints. Given the likely biological implications of 3’-seq signal inside of gene coding regions and other genomic regions, and the mixed accuracy of available annotations, it is important to ensure that 3’-end data is comprehensively assessed in annotation agnostic ways.

While providing a cohesive and statistically grounded approach to 3’-seq analysis, PIPETS is also designed to be adjustable to account for diffierences between data sets. We invite users to analyze their data with diffierent strictness levels of PIPETS to identify a global threshold that they are comfortable assigning biological significance to. In the case of data with low total read depth, we recommend running PIPETS with a lax strictness (increasing threshAdjust) to ensure that low read coverage values are not ignored. Conversely, data with high total read depth should be analyzed with a more strict (decreasing threshAdjust) version of PIPETS. In both circumstances, the ideal outcome is the identification of all large read coverage positions while also ensuring that low read coverage positions that derive from biologically significant events are not missed. However, even with the best possible set of parameters, PIPETS will likely fail to identify some biologically relevant 3’ sites or under analyze certain proportions of the data.

However, unlike previous 3’-seq analysis approaches, PIPETS does not arbitrarily exclude annotated coding regions from analysis, allowing users to fully utilize their data to inform biological conclusions.

In conclusion, PIPETS provides a novel, easily accessible, platform with which to explore 3’-seq data from diffierent organisms for the entire field. Even with comprehensive data sets and annotations from well-studied model organisms like *E. coli* and *B. subtilis*, there is still a fundamental lack of study across data sets from diffierent species. As there are no genome wide- ground truth data from orthogonal methods available, it is difficult to assess the validity of 3’- seq data generally. However, with its statistically informed significance measures, PIPETS is able to more confidentially parse signal from noise in 3’-seq data, and identify potentially biologically significant results in genome regions that have previously been un-analyzed. Furthermore, the parameterization of PIPETS enables users to explore their data, potentially adding whatever post-processing filters they see fit, while at the same time providing a common framework that can applied across data sets of diverse sizes from diverse organisms.

## Availability and requirements

**Project Name:** PIPETS: Poisson Identification of Peaks from Term-Seq data

**Project home page:** https://bioconductor.org/packages/release/bioc/html/PIPETS.html or https://github.com/qfurumo/PIPETS

**Operating system(s):** Platform independent

**Programming language:** R **Other requirements:** R >4.4.0 **License:** GPL-3

Any restrictions to use by non-academics: none.

## Abbreviations

PIPETS: (**P**oisson **I**dentification of **PE**aks from **T**erm-**S**eq data) UTR: **U**n**T**ranslated **R**egion

## Declarations

Ethics approval and consent to participate: N/A Consent for publication N/A

## Availability of data and materials

This work only utilizes existing publicly available datasets and annotation files including: Bioproject PRJEB12568 https://www.ncbi.nlm.nih.gov/bioproject/PRJEB12568), with genome annotation AL009126 (https://www.ncbi.nlm.nih.gov/nuccore/AL009126) and NCBI Short Reach Archive SRP136114 (https://www.ncbi.nlm.nih.gov/sra/?term=SRP136114), with genome annotation refseq: NC_003028.3 (https://www.ncbi.nlm.nih.gov/nuccore/194172857). Software availability is described above.

## Author Contributions Statement

Q.F. and M.M. conceptualized the software, Q.F. developed the software and published it, Q.F. and M.M analyzed the results. Q.F. and M.M. wrote and reviewed the manuscript.

## Supporting information

Additional Datafile 2

Additional Datafile 3

Additional Datafile 1

## Acknowledgements

We would like to thank Irem Ozkan and Leo Ahumada Manuel for their invaluable assistance as early testers of PIPETS. We would also like to thank the Bioconductor team for their support during the submission process.

## Additional Files

Supplementary Figures: Furumo_Supplementary_Figs.pdf, pdf file, Demonstration of Results from PIPETS parameter testing demonstrating insensitivity to minor parameter adjustments.

Supplementary Table 1: SupplementalTable1.csv, csv file, Run time analysis of PIPETS along coupled with diffierent threshAdjust cutoffis for each data set.

Additional Datafile 1: Warrier_PIPETS_Results.csv, csv file, all PIPETS predicted peaks (default parameters) from Warrier et. al. dataset described.

Additional Datafile 2: Dar_PIPETS_Results_threshAdjust075.csv, csv file, all PIPETS (threshAdjust=0.75) predicted peaks from Dar et al. dataset.

Additional Datafile 3: Dar_PIPETS_Results_threshAdjust085.csv, csv file, all PIPETS (threshAdjust=0.85) predicted peaks from Dar et al. dataset.

