## Supplemental Figures for "PIPETS: A statistically informed, gene-annotation agnostic analysis method to study bacterial termination using 3’-end sequencing"

**Furumo and Meyer: PIPETS (Poisson Identification of Peaks in Term-Seq data).**
**Supplemental Figures:**

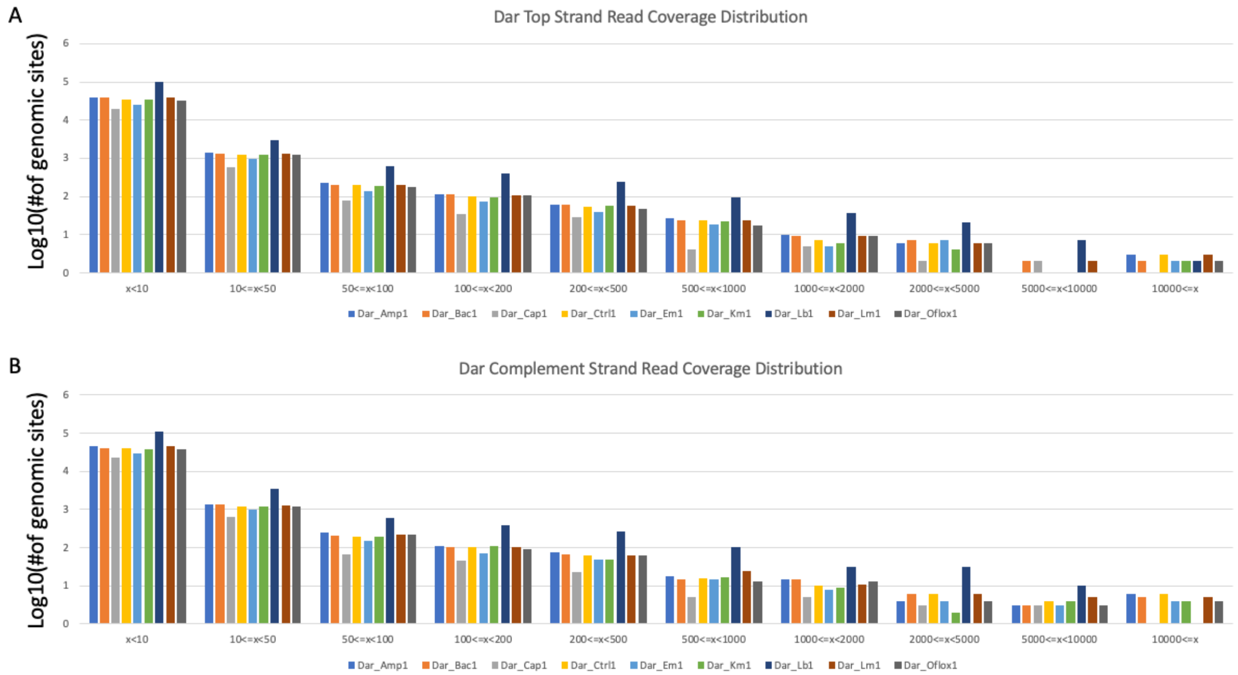


**Supplemental Figure 1: Distribution of read quality passing read coverage for positions across representative Dar data sets.** For both the top (A) and complement (B) strands of representative data sets from the Dar data (Dar *et al.* 2016), a majority of the genomic coordinates have read quality score passing read coverage of less than 10. Counts of genomic sites on the Y axis are on a Log10 scale.

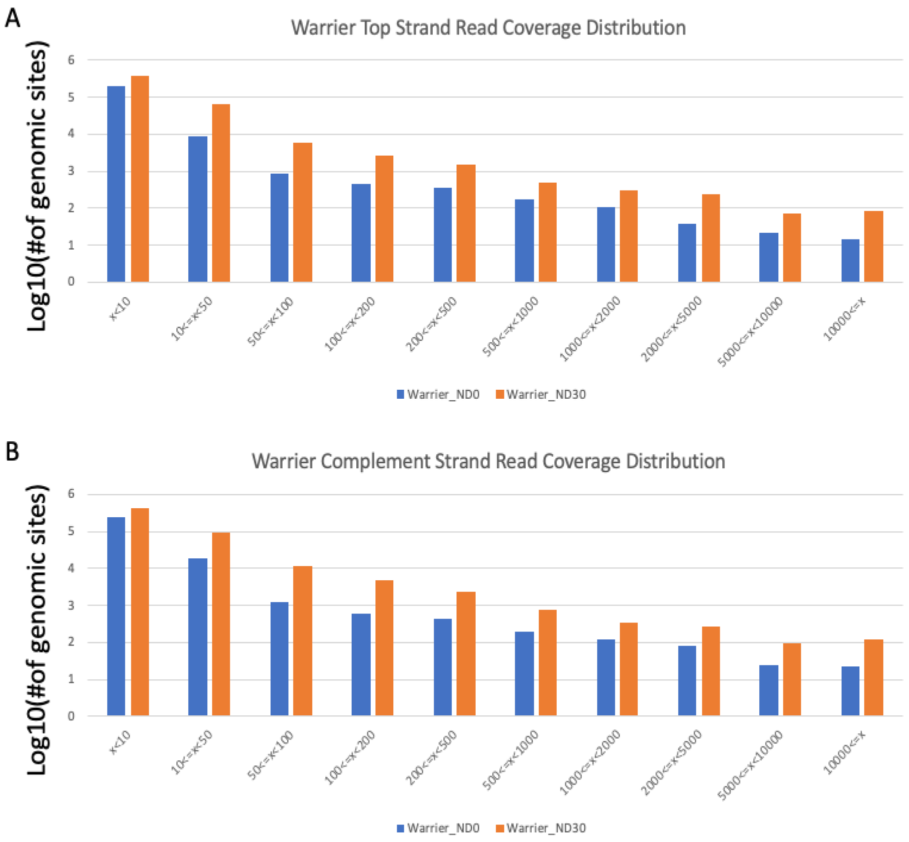


**Supplemental Figure 2: Distribution of read quality passing read coverage for positions across the Warrier data sets.** For both the top (A) and complement (B) strands of the data sets from the Warrier data (Warrier *et al.* 2018), a majority of the genomic coordinates have read quality score passing read coverage of less than 10. Counts of genomic sites on the Y axis are on a Log10 scale.

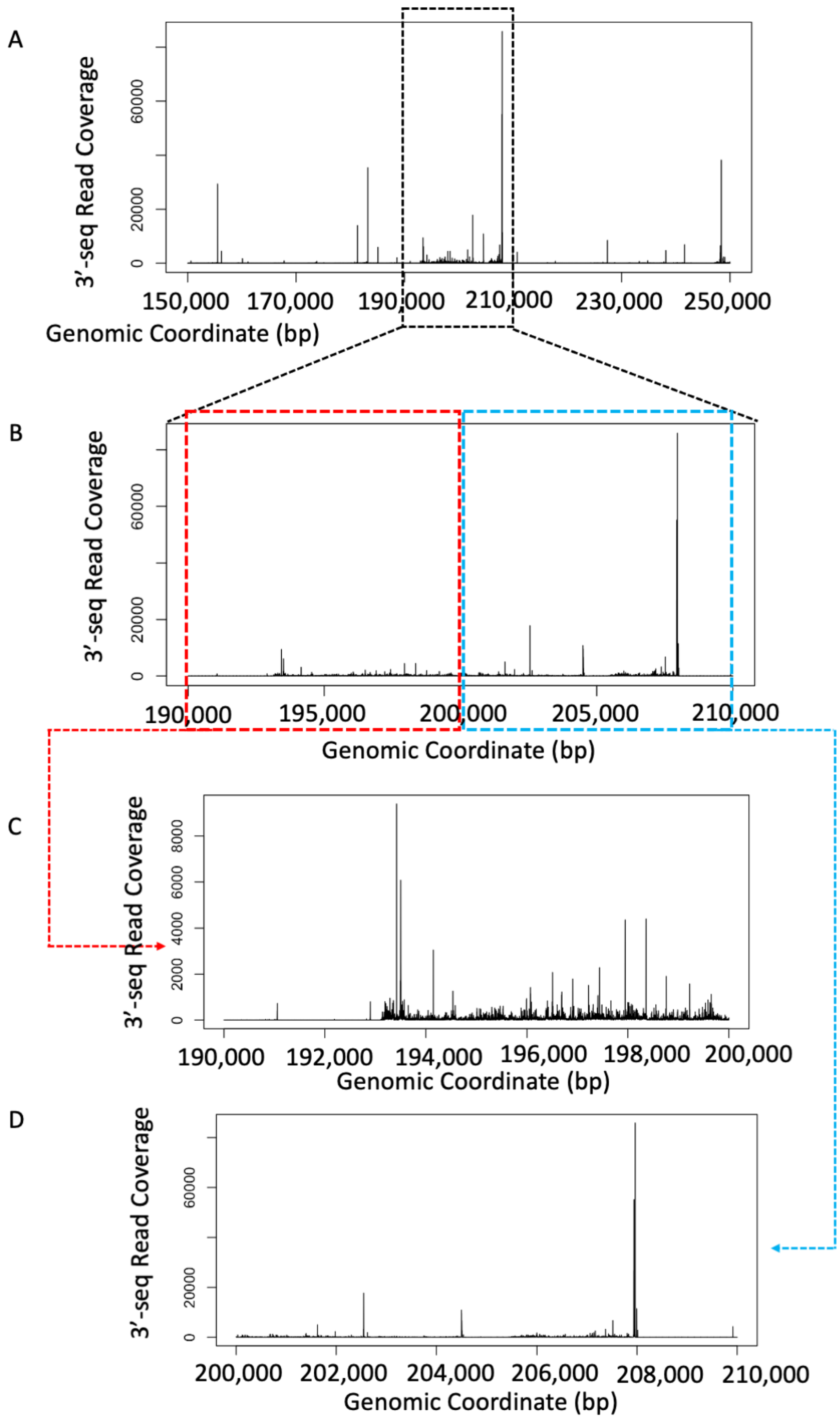


**Supplemental Figure 3: Prevalent low-level signal inside of likely gene coding region provides necessity for condensing of results.** (A,B) Read quality score passing read coverage levels from a 100,000 bp region from the Warrier_ND0 complement strand reads shows a likely pattern for a gene coding region. When this potential gene coding region is further broken down into two sections, two distinct patterns of 3'-seq read coverage emerge. (C) In the first half of the region, the read coverage is an order of magnitude lower than that of the total gene coding region but there are still many coordinates with read coverage throughout. Between positions 190,000 – 200,000, PIPETS identified 20 condensed peaks with significant 3’-seq signal. These 20 peaks contained 40 individual genomic positions with significant 3’-seq signal. While the coordinate with the highest read coverage for these peaks is reported as the primary 3’-seq signal for that peak, the total width of each peak is maintained for further study. (D) A small subset of the second half of the region dominates the 3’-seq read coverage, with 3’-seq read coverage several orders of magnitude higher than most of the signal. Between positions 200,000 – 210,000, PIPETS identified 28 condensed peaks with significant 3’-seq signal. These 28 peaks contained 127 individual genomic positions with significant 3’-seq signal. While this region spans 10,000 bp, the 85 bp range of 207926-208011 contains a large proportion of the total signal for the region.

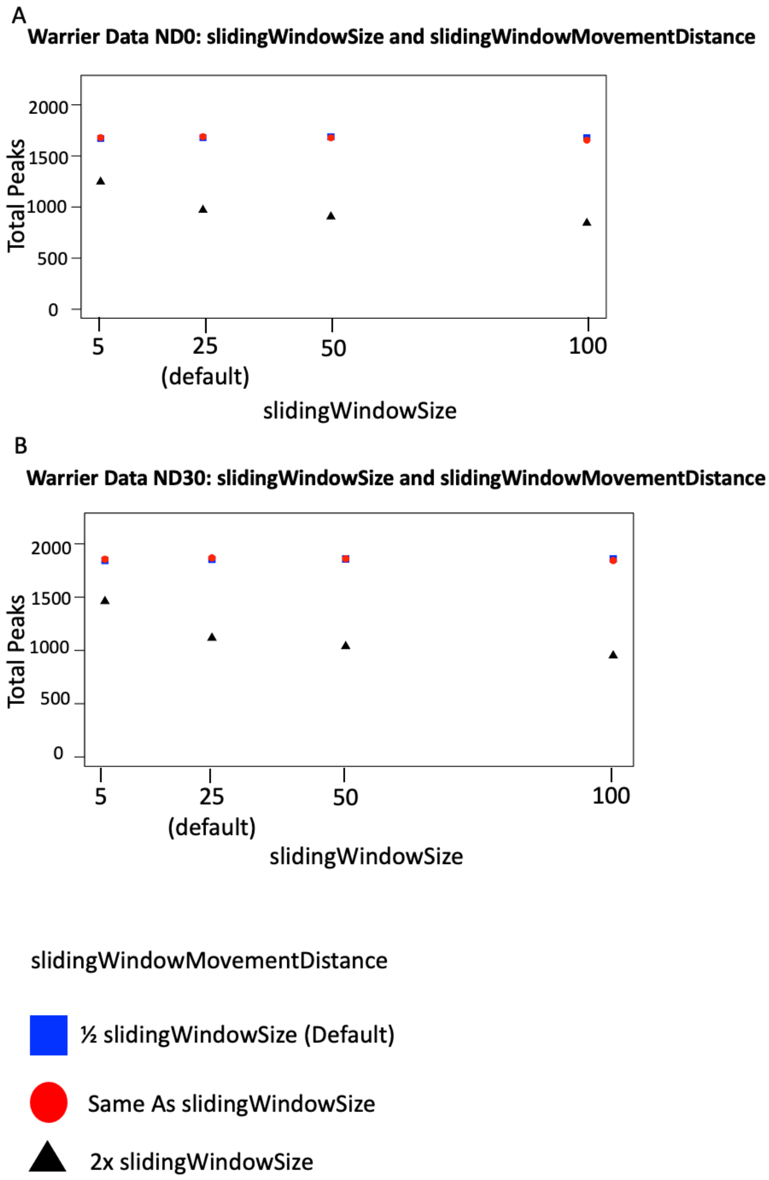


**Supplemental Figure 4: Testing robustness of PIPETS analysis to changes in slidingWindowSize and slidingWindowMovementDistance.** By default, slidingWindowSize is set to 25, which establishes a sliding window that extends 25bp upstream and downstream of a given coordinate, resulting in a window of total size 51 bp. slidingWindowMovementDistance is set to half, rounded-down, of that value (25bp) by default so that almost every position in the data is tested in two different windows. When testing differing slidingWindowSize values along with different proportional values of slidingWindowMovementDistance (1/2 x, x, 2x) on the Warrier ND0 (1A) and ND30 (1B) data sets, we noted that there was no dramatic change in the number of peaks identified by PIPETS, with the exception of when slidingWindowMovementDistance was so much larger than the slidingWindowSize that PIPETS was effectively skipping over genomic coordinates, and as such would not be a viable parameter set for study.

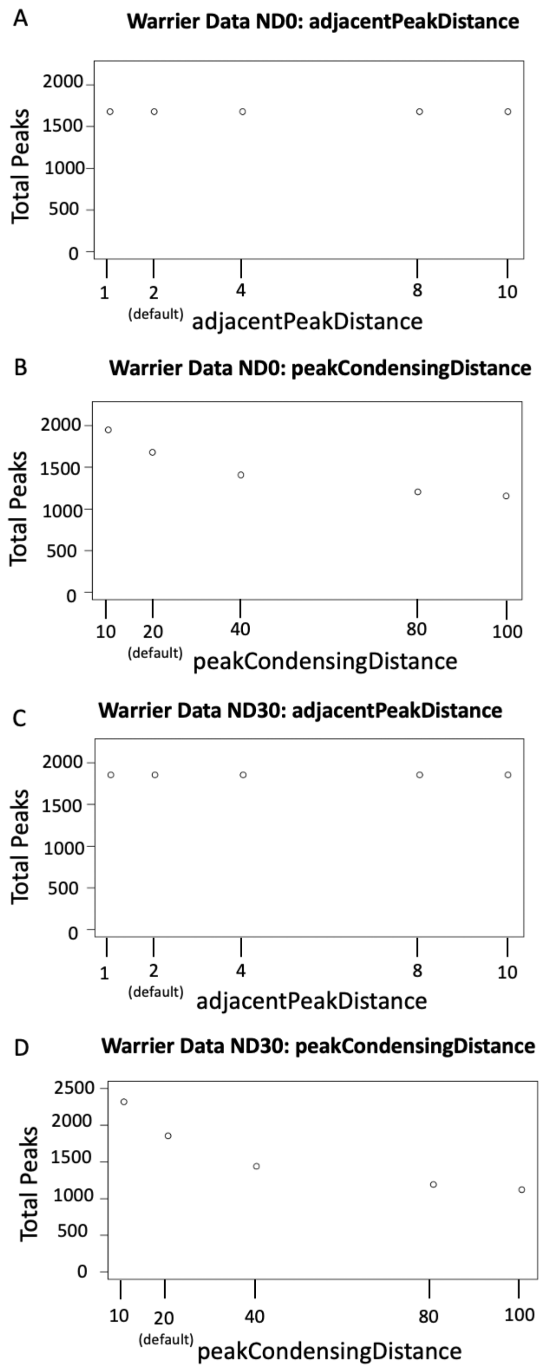


**Supplemental Figure 5: Testing robustness of PIPETS analysis to changes in adjacentPeakDistance and peakCondensingDistance.** PIPETS performs two results condensing steps to make results more palatable. First, it takes significant results that are located very within 2 bp of each other by default (adjacentPeakDistance =2) in the genome and condenses them into a singular 3'-seq "peaks", as they are unlikely to be individually significant events. Second, PIPETS takes the list of significant peaks and combines those that are withing 20bp of each other by default (peakCondensingDistance) as the likelihood of strong individual peaks at that proximity is low. In testing different values for both parameters, we noted that large changes in adjacentPeakDistance had no effect on the total number of peaks identified for either the Warrier ND0 (2A) or ND30 (2C). For peakCondensingDistance, we noted that while there were some larger differences in the total number of peaks for very high values, there was not much change in total peaks found in the ranges close to the default (20 nt) for the Warrier ND0 (2B) and ND30 (2D) data.
